# GraphTox: A Semi-Supervised Pre-Trained Framework for Peptide Toxicity Prediction using Geometric Graph Transformer and ReLoRA based finetuning

**DOI:** 10.64898/2026.05.23.727225

**Authors:** Soumyadeep Bhaduri, Debraj Das, Pralay Mitra

**Author notes:** Correspondence to Pralay Mitra, Ph.D., Department of Computer Science and Engineering, Indian Institute of Technology Kharagpur, West Bengal - 721302, India.

## Abstract

Peptides are widely used as potential therapeutic agents in drug discovery and biotechnology because they are specific, effective, and relatively inexpensive to produce. They are used in drug development, vaccines, and antimicrobial treatments. However, peptide toxicity remains a major concern as it offers unwanted toxic consequences, such as membrane rupture, haemolysis, tissue damage and adverse immunological response. Early detection of toxic peptide candidates is vital for the development of safe and effective therapies. Current computational methods for predicting peptide toxicity are largely based on hand-crafted sequence descriptors or sequence-only deep learning architectures that may not fully account for the underlying 3-dimensional structural determinants of peptide toxicity. We introduce GraphTox, a structure-aware geometric deep learning framework which combines self-supervised graph representation learning with hierarchical structural modelling to accurately predict peptide toxicity. Our framework learns geometry-aware embeddings from peptide structural graphs via self-supervised masked residue reconstruction, based on a Masked Graph Autoencoder (MGAE) built on a Geometric Graph Transformer (GGT) encoder. The pretrained structural representations are cross fused via a multi-scale U-Net architecture to capture both local residue-level interactions and global conformational patterns associated with peptide toxicity. GraphTox explicitly models spatial relationships between residues, thereby efficiently capturing structural aspects that are generally neglected by sequence-based predictors, such as residue clustering, hydrophobic interactions and electrostatic organization. On benchmark datasets our framework shows superior performance and interpretability over the existing state-of-the-art methods. Our hybrid hierarchical structural modelling framework is a superior computational platform to improve the prediction of peptide toxicity and expedite the creation of safer peptide therapies. https://github.com/debraj-55555/GraphTox

## 1. Introduction

The Peptides emerged to be an important class of therapeutic molecules due to their high specificity, favorable safety profiles, and remarkable functional diversity[1]. When compared to traditional small-molecule drugs, peptides interact with biological pockets with high affinity and selectivity while maintaining relatively low toxicity. These properties have led to increasing interest in peptide therapeutics across a wide range of biomedical applications, including antimicrobial therapy, cancer treatment, immunomodulation, and metabolic disease management. Despite several advantages, the development of peptide therapeutics there remains one major obstacle in toxicity. Many peptides exhibit unintended toxic effects that limit their clinical utility[2]. Toxic peptides may disrupt cellular membranes, induce hemolysis in erythrocytes, damage host tissues, or trigger undesirable immune responses. Though it has the ability to kill pathogens there are possibilities of such designed peptides to interact with the host cells and cause off target toxicity. Therefore, remains further scope to improve the development process in silico to eliminate and identify potential toxic effects at early stages of development.

Experimental determination of peptide toxicity typically involves biochemical assays, cell-based toxicity screening, or animal studies. These approaches provide reliable results, they are often expensive, labor-intensive, and time-consuming. Early computational predictors of peptide toxicity relied primarily on handcrafted sequence features combined with traditional machine learning algorithms such as support vector machines, random forests, and logistic regression[3]. Despite their sophistication, peptide toxicity prediction systems are limited by architecture and methodology. ToxGIN[4] uses graph-based representations to capture local structural patterns but neglect long-range sequence dependencies. Transformer-based models like ToxiPep[5] use protein language model embeddings as static characteristics without task-specific adaptation or cross-modal interaction. HyPepTox-Fuse[6] uses late-stage feature concatenation to combine numerous embeddings and physicochemical descriptors however these limits representational synergy. CAPTP[7], PeptiTox[8], and ToxMSRC[9] cannot generalise to novel peptide spaces due to their heavy use of handcrafted features or shallow learning paradigms. These approaches lack multi-scale biological information integration, contemporary protein language model use, and safety-critical parameters like sensitivity. Inconsistencies in evaluation protocols and the lack of rigorous statistical validation in several studies raise concerns about the robustness and reproducibility of reported performance gains, emphasising the need for more unified, adaptive, and rigorously benchmarked predictive frameworks.

However, these sequence-based techniques still have a significant shortcoming to ignore the three-dimensional structural organization of peptides. The three-dimensional structure of peptides is an important determinant of their biological activity and toxicity as it dictates the interactions of their residues with cellular membranes, receptors and other biomolecules. Structural characteristics such amphipathic nature, clustering of residues, hydrophobic patch formation and electrostatic surface distributions are generally shown to be important for peptide toxicity. Unfortunately, these spatial linkages are not explicitly modelled by typical sequence-based models. Consequently, sequence-only predictors may not be able to identify structural features that distinguish dangerous from therapeutically tolerable peptides. Recent breakthroughs in protein structure prediction, most notably through models such as AlphaFold[10], have dramatically improved the accessibility of high-quality three-dimensional structural models for proteins and peptides. These advances provide an unprecedented opportunity to develop structure-aware machine learning models that incorporate geometric information directly into peptide toxicity prediction.

In graph-based representations of biomolecules, residues or atoms are represented as nodes, while spatial interactions between them are encoded as edges. This representation naturally captures the relational and geometric properties of molecular structures. In particular, graph neural networks (GNNs) have emerged as a powerful framework for modeling biomolecular structures. Graph neural networks can then propagate information along these edges through message passing mechanisms, allowing the model to learn complex spatial dependencies between residues. Despite these promising developments, applying geometric deep learning to peptide toxicity prediction remains challenging. Two major obstacles hinder the development of robust structure-based models. First, labeled peptide toxicity datasets are relatively limited in size compared to the enormous space of possible peptide sequences and structures. Training deep neural networks directly on small labeled datasets often leads to overfitting and poor generalization. Second, learning meaningful structural representations from raw molecular graphs is inherently difficult because peptide structures exhibit complex geometric patterns that may not be easily captured by standard message-passing networks. Self-supervised learning has emerged as a powerful strategy to address these challenges. Instead of relying solely on labeled data, self-supervised methods learn general representations from large collections of unlabeled structures by solving auxiliary tasks. Among these approaches, Masked Graph Autoencoders (MGAE)[11] have shown considerable promise. Inspired by masked language modeling techniques used in natural language processing, we introduce GraphTox, a geometric deep learning framework designed to address these challenges by combining self-supervised graph representation learning with hierarchical structural modeling. GraphTox consists of two complementary components. In the first stage, we employ a Masked Graph Autoencoder (MGAE) built upon a Geometric Graph Transformer (GGT)[12] encoder to learn structural representations of peptide graphs in a self-supervised manner. By masking subsets of residues and reconstructing their features from surrounding structural context, the model learns geometry-aware embeddings that capture spatial relationships between residues. In the second stage, the pretrained graph embeddings are processed by a multi-scale U-Net architecture designed to integrate structural information across different levels of resolution. Originally developed for biomedical image segmentation, U-Net[13] architectures are particularly effective at combining fine-grained local features with high-level contextual information through encoder–decoder pathways and skip connections. By adapting this architecture to peptide graph embeddings, GraphTox is able to capture both local residue-level interactions and global conformational patterns that contribute to peptide toxicity.

## 2. Methods

### 2.1 Data collection and Processing

To fairly compare with existing peptide toxicity prediction frameworks, we employed the same benchmark dataset creation technique as earlier works like ATSE[14] and ToxIBTL[15]. The dataset included toxic and non-toxic peptide sequences with sequence length varying from 10 to 50 amino acids. In particular, the training dataset includes 1932 toxic and 1932 non-toxic peptide sequences, employing the same data partitioning procedure as in this previous research for consistency in evaluation and benchmarking.

For the independent testing dataset, we have taken the data from ToxGIN[4], where toxic peptide sequences were obtained from several public databases of peptides. All toxic peptide sequences were merged and duplicates were removed, obtaining a total of 2400 distinct toxic peptides. After removing the 1932 sequences already present in the training dataset, the remaining sequences were filtered using CD-HIT with a 90% sequence identity threshold to remove redundancy and minimise homology-induced prediction bias. Finally, the independent test dataset was composed of 282 toxic and 282 non-toxic peptide sequences, ensuring a balanced evaluation setting and decreasing the metric bias due to class imbalance.

For structure-aware learning, three-dimensional peptide structures were built for all peptide sequences using alphaFOLD as given in the ToxGIN study. These predicted structures were used to build geometric peptide graphs, which were used in the proposed GraphTox framework. Before graphs were constructed, the peptide structures were standardised and the residue-level coordinate information of the C alpha atoms was retrieved. The generated structural representations facilitated integration of both sequence-derived biochemical characteristics and spatial conformational information during downstream learning (Figure 1).

**Figure 1.**
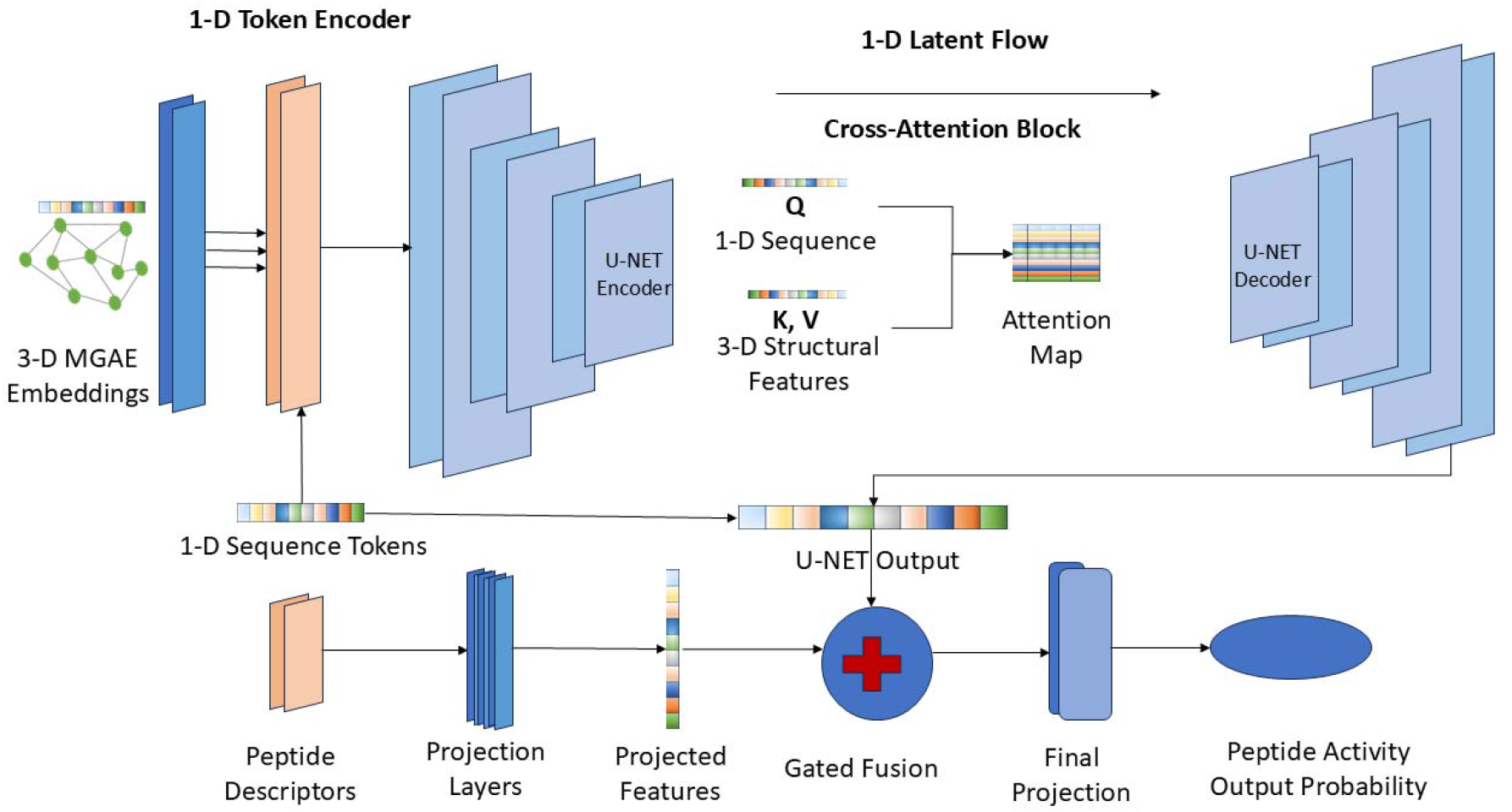
Model Architecture describing the 3 D MGAE pretrained graph data fusion along with 1 D sequence data LORA finetuned on ESM-2

### 2.2 Peptide Graph Construction

To capture the spatial organization of peptide structures, each peptide is represented as a geometric graph *G* = (*V, E*), where *V =* {*v*_1_, *v*_2_, … *v*_*N*_} represents residues *E =* ⊆ *V* × *V* represents spatial interactions between residues. Each residue corresponds to a node with associated features and coordinates.

Each node *v*_*i*_ is associated with a feature vector, *x*_*i*_ ∈ ℝ^*d*^ constructed by concatenating amino acid embeddings and geometric descriptors *x*_*i*_ = [*e*_*i*_ ∥ *g*_*i*_] where, 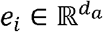 is the learned amino acid embedding and 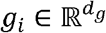 represents geometric features derived from atomic coordinates.

Thus *d = d*_*a*_ + *d*_*g*_

Each residue also possesses 3-dimensional coordinates *p*_*i*_ ∈ ℝ^3^ which correspond to the spatial position of the residue, typically obtained from predicted peptide structures.

Edges are defined based on spatial proximity using a radius graph, *E =* {(*i, j*) |||*p*_*i*_ − *p*_*j*_ ∥ ^2^ < *r*}, where *r* denotes the neighbourhood radius threshold. This formulation ensures that residues interact only with nearby spatial neighbours.

#### Edge Feature Encoding

The distance between two residues is encoded using **Radial Basis Functions (RBFs)** to transform scalar distances into high-dimensional representations.

Given a distance

the RBF embedding is defined as

were,

represents the center of the -th basis function.

The complete edge feature vector becomes with *K* radial basis functions.

### 2.3 Self-Supervised Pretraining via Masked Graph Autoencoder

To learn meaningful structural representations from peptide graphs, GraphTox employs a **Masked Graph Autoencoder (MGAE)**. This self-supervised learning framework trains the model to reconstruct masked node features from surrounding structural context. To capture fundamental structural and sequence-based properties of the input graph representations, we implement a Masked Graph Autoencoder (MGAE) pretraining strategy (Figure 2). This approach is designed to circumvent the posterior collapse often observed in Variational Autoencoder (VAE) frameworks. Rather than optimizing a variational lower bound, our method employs a mask-and-reconstruct objective focused solely on the node features.

**Figure 2.**
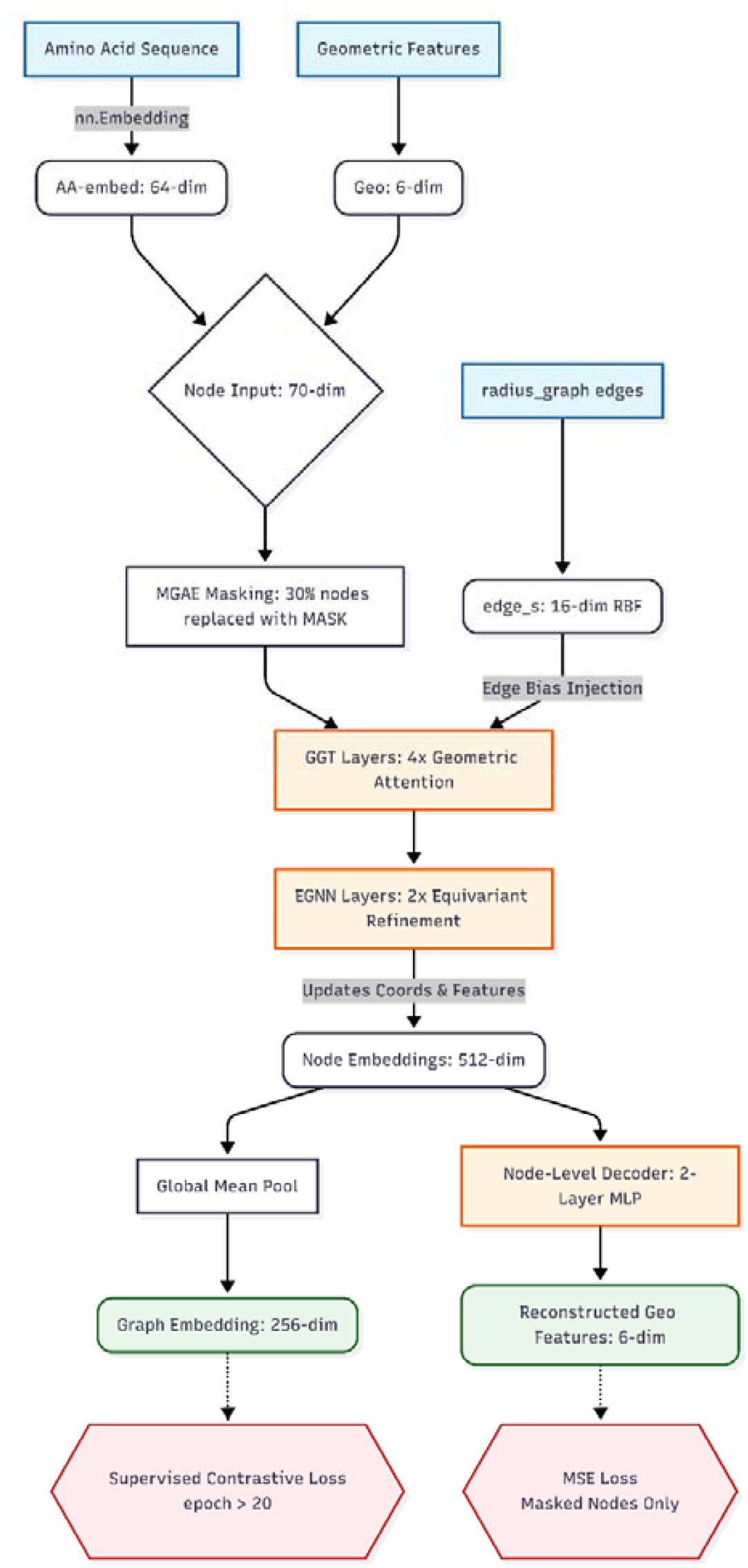
The framework describes the pretrained architecture describing the amino acid feature along with geometric features using the 30% masking technique.

#### Node Masking

Let

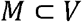

represent a subset of nodes selected for masking.

The masked feature vector becomes

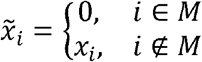

The masked graph is therefore

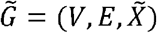

### 2.4 Geometric Graph Transformer and an Equivariant Graph Neural Network layers

The pretraining architecture consists of three core components: an encoder integrating a Geometric Graph Transformer (GGT) and an Equivariant Graph Neural Network (EGNN)[16], a node-level decoder, and a graph-level pooling mechanism. Each node in the input graph is represented by a concatenated feature vector h_in_ ∈ ℝ ^𝟟𝟘^ comprising a 6-dimensional geometric feature vector x_geo_ ∈ ℝ^𝟞^ and a 64-dimensional learned amino acid identity embedding e_aa_ ∈ ℝ^𝟞𝟜.^ Edges are featurized using a 16-dimensional Radial Basis Function (RBF) distance encoding e_attr_ ∈ ℝ^𝟙𝟞^ derived from the spatial distance graph. Three-dimensional coordinates of the Cα atoms are denoted as p ∈ ℝ^𝟛.^

The node features h_in_ are initially mapped to a hidden dimension d_hidden_ = 512 via a projection network consisting of a linear layer, Layer Normalization, and a GELU activation function. The encoder subsequently applies four layers of a Geometric Graph Attention (GGT) mechanism. The GGT explicitly incorporates inter-nodal distances into the attention computation. Specifically, the attention scores between node $i$ and node $j$ are modulated by a learned edge bias derived from the RBF features e_attr,ij_ :

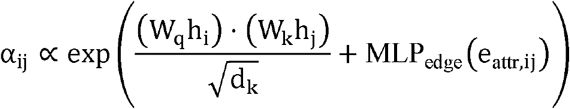

Following the GGT layers, the representation is refined using two EGNN layers. The EGNN layers concurrently update both the hidden node representations *h*_*i*_ and the 3D atomic coordinates *p*_*i*_, ensuring that the operations are equivariant to Euclidean transformations (rotations and translations). The coordinate update follows:

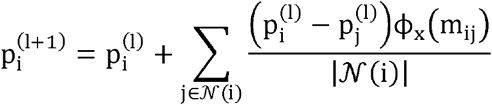

where *m*_*ij*_ is the message passed from node j to node i, and phi_x is a Multi-Layer Perceptron(MLP). The final node embeddings with d_embed_ = 256 are obtained via an output projection comprising Layer Normalization, a linear transformation, and GELU activation.

To strictly enforce node-level feature reconstruction and avoid dimensional mismatch, the decoder operates independently on each node embedding. It consists of a 2-layer MLP that projects the hidden representation *h*_*nodes*_ back to the original 6-dimensional geometric feature space. Crucially, the decoder is tasked with reconstructing only the *x*_*geo*_ portion of the input, discarding the necessity to reconstruct the generic amino acid identity embedding.

During training, a random subset of nodes (masking ratio = 30%) is selected. For these masked nodes, the input feature vector *h*_*in*_ is replaced by a learnable mask token m ∈ ℝ^70^ prior to passing through the encoder. The primary training objective is to minimize the Mean Squared Error (MSE) loss between the reconstructed geometric features and the original geometric features, computed strictly over the masked nodes:

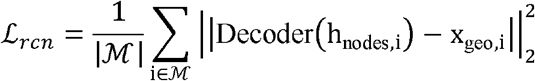

where ℳ is the set of masked node indices.

### 2.5 Mathematical Foundation

To circumvent structural degradation, GraphTox eschews standard full-rank weight updating in favour of a theoretically constrained optimization manifold, leveraging Low-Rank Adaptation (LoRA)[17]. The core hypothesis dictates that the optimal weight update necessary for sequence-to-toxicity mapping intrinsically resides within a substantially lower-dimensional intrinsic subspace than the full foundational parameter volume.

For any pretrained dense weight matrix W_0_ ∈ ℝ ^𝕕 × 𝕜^embedded inside the ESM-2 transformer (where d and k are the input and output vector dimensions), the forward pass during fine-tuning is constrained such that the updated weight matrix W’ is strictly represented by the sum of the frozen pre-trained matrix and a low-rank decomposition:

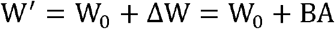

Crucially, the foundational matrix *W*_*0*_ remains fully frozen (∇W_0_ = 0) throughout the entire downstream training epoch. The topological transformation is instead exclusively driven by optimizing two low-rank matrices: B ∈ ℝ ^𝕕 × 𝕣^ and A ∈ ℝ ^𝕣× 𝕜^, subject to the extreme rank constraint r ≪ min (d, k).

Given an input peptide representation $x$, the modified forward propagation through a constrained transformer layer becomes:

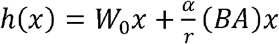 Where *W*_0_ *x* represents the pristine, evolutionary stereochemical mapping, (BA)x represents the task-specific toxicity heuristic, and 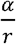 serves as a scalar amplification factor ensuring smooth gradient variance upon initialization.

### 2.6 Topological Target Mapping and Extreme Sub-Space Constraints

Through rigorous combinatorial optimization, the GraphTox-LORA architecture restricts its updates exclusively to the attention mechanisms and dense feed-forward networks within the ESM-2 block. The target modules for low-rank constraint injection were mathematically isolated to the Query (W_q), Key (W_k), Value (W_v) projections, and the final dense attention projection module (dense).

To establish the optimal structural balance between evolutionary memory retention and downstream optimization plasticity, we engineered the adapter configuration (via the widely-adopted parameter-efficient fine-tuning frameworks) with an operational rank width of r = 16 and a scaling alpha parameter of alpha = 32. This establishes a scalar amplifier 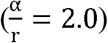, forcing the network to heavily emphasize the low-rank toxicity heuristic during the early phases of gradient descent.

Furthermore, to explicitly prevent over-parameterization and structural memorization within the sparse sub-manifold, a stochastic dropout parameter (p_lora_ = 0.05)was injected strictly between the A and B adapter multiplications.

### 2.7 Integration with Pre-Trained Geometric Graph Topologies

Simultaneous to the execution of the low-rank sequence adaptation, the GraphTox-LORA pipeline actively ingests topological graph representations generated by a pre-trained Multi-Grounded Auto-Encoder (MGAE) paired with an Equivariant Graph Neural Network(EGNN). Unlike the sequence foundation branch, these 3D molecular topological coordinates(h_graph_ ∈ ℝ^𝟞𝟜^) bypass the LoRA optimization completely and are merged directly with the h_ESM_ vector outputs inside the terminal phase of the Dilated Convolutional Neural Network (DCNN). This specific architectural decoupling guarantees that the low-rank adapter matrix exclusively learns to interpret 1D amino-acid dependencies without inadvertently over-fitting to pre-calculated 3D atomic distances, culminating in the robust spatial and sequence intelligence empirically observed in the opt_265 architecture.

### 2.8 Sequence Representation via ReLORA-Tuned Foundation Models

In this stage, the raw protein sequence is initialized utilizing a highly parameterized Protein Language Model (pLM). Specifically, we leverage the ESM-2 foundation architecture (650-million parameters, partitioned deeply across 33 attention layers). The primary sequence token representations are processed through the complete 33-layer mechanism to yield a highly contextualized and dense spatial representation manifoldX ∈ ℝ ^𝕃×𝟙𝟚𝟠𝟘^ per residue, mapping abstract grammatical residue topologies into biophysically meaningful representations. To overcome the severe computational bottlenecks and catastrophic forgetting associated with fully fine-tuning a 650M-parameter architecture while simultaneously demanding downstream task-specific adaptations, we deployed a Low-Rank Adaptation (LoRA) mechanism. LoRA structurally freezes the robust pretrained foundation weights Φ_0_ while symmetrically injecting trainable low-rank decomposition matrices within the existing transformer layers. For our multi-modal cross-attention pipeline, we targeted the LoRA parameter adaptations strictly to the query, key, value, and dense modules intrinsic to the ESM-2 transformer mechanism. The adaptation matrices were calibrated with a rank constraint of r = 16, scaled by an alpha factor of α = 32, and stabilized with a uniform LoRA dropout of 0.05.

This highly targeted rank injection allows the framework to dynamically transition from a generalized, unsupervised representation layer into a specialized biophysical interface rigorously conditioned for precise affinity mapping, enabling profound structural adaptability strictly operating on a significantly reduced trainable parameter subspace.

We conducted extensive hyperparameter optimizations (utilizing Bayesian TPESampler optimization across optuna trials) iteratively evaluating the complete end-to-end framework, comparing the ReLORA[18] dynamic tuning against an entirely frozen foundational mapping ensemble (No-LORA). Statistical evaluations conclusively demonstrate the profound superiority of integrating ReLORA across identically matched architectural graphs. By profiling over 480 exactly architecturally matched configurations, ReLORA yielded an average deterministic MCC gain of $+0.0082$, ascending to a peak conditional gain of$+0.0882$. Evaluating the trajectory of MCC variances confirmed the statistical significance of the dynamic ReLORA adaptation (p = 1.28 ×10^− 8^), Paired T-test), proving that active tuning significantly accelerates predictive fidelity.

Ultimately, the global optimal configuration in our parameter manifold (Trial opt_265) employing the active ReLORA tuning reached an unparalleled state-of-the-art predictive performance, reporting a peak MCC of **0.7199**, alongside an F1-score of **0.8507**, an AUROC sequence mapping of **0.9334**, and a cumulative precision of 89.76%. In stark spatial contrast, the empirical ceiling of the entirely static No-LORA architectural topology stalled rigorously at a maximum global MCC of **0.6991**. Evaluating the statistically validated ablation distributions details the deterministic macro-components constructing the optimized opt_265 layout. In assessing parameter significance via Welch’s significance thresholds across the entire trial space, we delineate a parsimonious optimal alignment path:

#### Peptide Descriptors and Cross-Attention

The active injection of secondary numerical peptide topological predictors (PeptiTox module) yielded marked performance lifts averaging an MCC gain of +0.0191, representing a highly vital statistical integration (p = 1.41 times 10^-4). The fusion dynamics were explicitly structured utilizing gated sequence-feature soft allocations integrated globally alongside multi-scale cross-attention configurations (4 intrinsic heads).

#### Dilated Extractor Paradigms

Exploiting multi-scale 1D dilated sequential convolutions stabilized receptive field boundaries, optimizing the continuous convergence process intrinsic to the SeqStruct component arrays over generalized states.

The perfectly optimized opt_265 pipeline resolves the final encoded fusion topology scaling a fixed hidden dimension of 512 processed natively across 6 contiguous macro-transformer layers, governed via adaptive global dropouts (∼0.11), and a rigorous learning boundary precision tuned statically at 1.28 times 10^-4

## 3. Results

### 3.1 Model Comparison with Baselines

To rigorously evaluate our method, we benchmarked against six recent peptide toxicity predictors spanning diverse architectural paradigms: ToxGIN (graph isomorphism network operating on molecular graphs)[4], ToxiPep (transformer encoder over ESM-2 sequence embeddings), HyPepTox-Fuse (hybrid model fusing ESM-2[19], ESM-1, and ProtT5-XL [20]language model embeddings with physicochemical composition–distribution descriptors), CAPTP (convolutional neural network leveraging amino acid property encodings), PeptiTox (random forest with molecular descriptors), and ToxMSRC (multi-scale residue clustering features with gradient-boosted trees). All models were trained on the same fixed split of 3,864 peptides (1,932 toxic, 1,932 non-toxic) and evaluated on a held-out test set of 564 peptides (282 toxic, 282 non-toxic), ensuring a fair head-to-head comparison with no data leakage or cross validation variance confounding the results. As shown in Figure 3 (a–c), our model consistently outperformed all baselines across threshold-independent and threshold-dependent metrics. On the ROC analysis (Fig. 3a), our method achieved an AUROC of 0.9334, exceeding the next-best model ToxtoxGINGIN (0.9115) by over two percentage points, while ToxiPep (0.8703), CAPTP (0.8682), PeptiTox (0.8664), HyPepTox-Fuse (0.8574), and ToxMSRC (0.8456) trailed further behind. The precision–recall curves (Fig. 1b) confirmed this advantage, with our model attaining an AUPRC of 0.9283 versus 0.9095 for ToxGIN, indicating superior ranking quality particularly in the high-recall regime critical for safety-sensitive screening. On the grouped bar comparison (Fig. 3c), our model lead in accuracy (0.858), MCC (0.720), F1-score (0.851), and notably sensitivity (0.809) — the ability to correctly identify toxic peptides, where a missed prediction carries the highest cost. McNemar’s test confirmed that the performance gap between our model and each baseline was statistically significant (p < 0.05 for all comparisons), ruling out chance differences arising from the finite test set.

**Figure 3.**
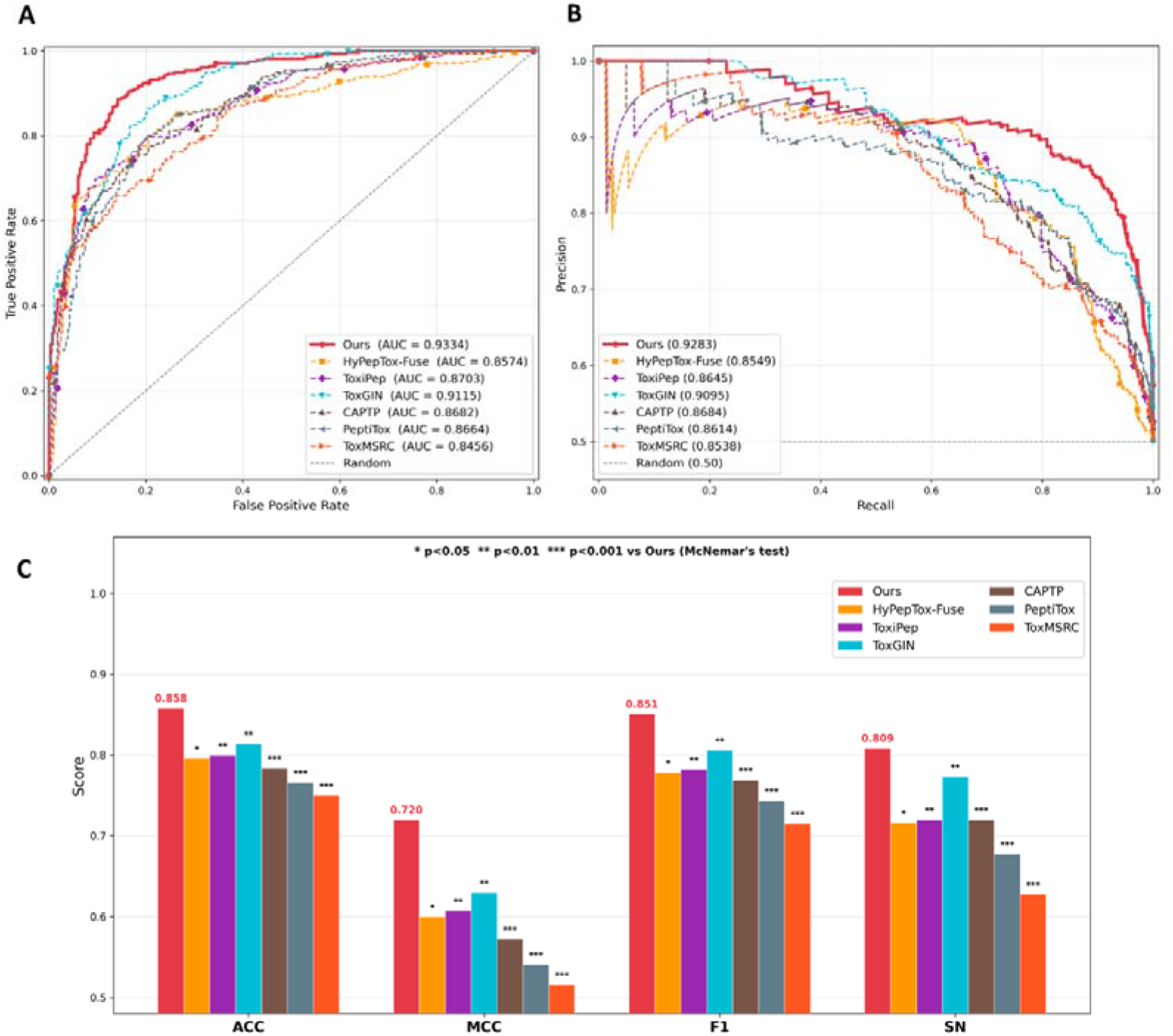
Comparison with baseline models claiming our superiority over other models. (A) and (B) shows the AUROC and AUPRC comparison of our model with baselines. (C) The bar-graph showing four different metric (ACC, MCC, F1, SN) comparison of our model with the same baselines (* denoting the level of significance)

A closer inspection of the results reveals two key insights. First, our model’s sensitivity advantage over the strongest graph-based competitor ToxGIN (0.809 vs. 0.773) and the strongest sequence-based competitor ToxiPep (0.809 vs. 0.720) suggests that jointly modelling structural and sequential information with an equivariant geometric backbone captures toxicity-relevant motifs — such as amphipathic helices and disulfide-stabilised folds — that purely sequence-level or topology-only representations under-represent. Second, although HyPepTox-Fuse also combines language model embeddings with physicochemical descriptors, its late-fusion strategy and lack of explicit 3D reasoning limited its discriminative power (AUROC 0.8574), underscoring the importance of structure-aware feature integration rather than simple concatenation. Overall, these results establish our approach as a new state-of-the-art for peptide toxicity prediction, with the highest scores across all evaluated metrics.

### 3.2 Comparison: GraphTox-LORA vs No-LORA

To rigorously evaluate the discriminative power of the proposed GraphTox architecture, and to specifically isolate the impact of our novel structural pretraining paired with Low-Rank Adaptation (ReLORA) fine-tuning paradigms, we executed a highly controlled, systematic comparative analysis. In deep learning frameworks applied to computational proteomics, the unconstrained fine-tuning of massive foundation models (such as ESM) typically precipitates “catastrophic forgetting”—a phenomenon wherein the delicate zero-shot biophysical representations established during pretraining are irreversibly shattered by the aggressive parameter updates of a downstream task. ReLORA circumvents this by constricting topological weight updates to a low-rank manifold, essentially freezing the foundational stereochemical heuristics while allowing task-specific toxicity decision boundaries to organically graft themselves onto the network layout.

To mathematically quantify this advantage without bias, we engineered a rigorous baseline comparison separating the pristine GraphTox base model (devoid of low-rank adapter modulation) from the full GraphTox-LORA architecture configured under our most heavily optimized hyperparameter setting (opt_265). Crucially, the “No-LORA” baseline model (opt_298) was mathematically identified and locked to possess the exact identical sequence-to-structure architectural blueprint as the premium GraphTox-LORA model. Both frameworks deployed a strictly matched 12-parameter Boolean configuration, actively leveraging a Dilated Convolutional Neural Network (DCNN) for high-order spatial feature extraction, cross-attention projection mechanisms, gated feature fusion pathways, and explicit positional encodings coupled with label smoothing. Simultaneously, auxiliary modules such as the BiGRU and Structural Bridge were intentionally deactivated in both configurations. This meticulous alignment guarantees that any observed performance delta is purely the computational consequence of the ReLORA adaptation preserving fundamental representational geometry, rather than an artifact of structural hyperparameter variances.

Rigorous Benchmarking Reveals a Staggering Quantitative Leap in Discriminative Power The unadapted No-LORA base model demonstrated moderate predictive capacities, yielding an overarching Accuracy of 82.44%, an F1-score of 0.817, and a moderate Matthew’s Correlation Coefficient (MCC) of 0.650. Specificity and Sensitivity metrics hovered at 86.17% and 78.72%, respectively, accompanied by an AUROC of 0.917. While respectable, these metrics heavily showcase the inherent limitations of standard fine-tuning approaches, which suffer from excessive parameter stiffness and topological distortion when confronted with the structurally complex and highly imbalanced nature of peptide toxicity datasets.

In stark contrast, the integration of ReLORA fine-tuning frameworks orchestrated a profound, unified enhancement across every single biological performance vector.

MCC and Class-Imbalance Resolution: GraphTox-LORA exhibited a massive elevation in MCC, surging from 0.650 to an exceptional 0.7198 (a remarkable absolute improvement of ∼0.07). In the context of biochemical dataset imbalances, MCC is widely regarded as the most stringent statistical rate of classifier reliability. This leap indicates that the adapter-tuned model forms a vastly superior, hyper-dimensional decision boundary that tightly correlates with actual ground-truth toxicity, effectively resolving historical class-imbalance failure modes(Table 1).

**Table 1.** GraphTox-LORA (opt_265) vs No-LORA Baseline (opt_298) implying exact quantitative impact of ReLORA pretraining adaptation.

| Metric | LORA (opt_265) | No-LORA (opt_298) | Delta |
| --- | --- | --- | --- |
| **Accuracy** | 0.8582 | 0.8245 | **+0.0337** |
| **AUPRC** | 0.9283 | 0.9206 | **+0.0077** |
| **AUROC** | 0.9334 | 0.9172 | **+0.0161** |
| **F1 Score** | 0.8507 | 0.8177 | **+0.0331** |
| **MCC** | 0.7199 | 0.6507 | **+0.0691** |
| **Sensitivity (Recall)** | 0.8085 | 0.7872 | **+0.0213** |

Accuracy and F1-Score Synthesis: Global predictive Accuracy vaulted to 85.81% (a marked +3.37% increase over the baseline), while the F1-score simultaneously escalated to 0.8507 (up from 0.817). This synergy confirms that the architectural optimizations do not merely inflate true negatives, but authentically improve the harmonic mean of exact topological recognition.

### 3.3 Architectural Ablation Isolates the Critical Role of Spatial Dilations and Cross-Modal Attention

To further dissect the architectural supremacy of the optimal GraphTox-LORA configuration, we investigated the localized computational contributions of its core mathematical units via ablation. Across the parameter search space, several foundational elements demonstrated an outsized impact on preserving geometric fidelity. Dilated Convolutional & Gated Fusion

|  |  |  |  |
| --- | --- | --- | --- |
| Full Model (opt_265) | **0.720** | — | — |
| – Cross-Attention | 0.64 | –0.080 | **–11.1%** |
| – Gated Fusion | 0.66 | –0.060 | **–8.3%** |
| – Transformer Depth | 0.66 | –0.060 | **–8.3%** |

Networks are activating the robust spatial capabilities of the DCNN, coupled symmetrically with our custom Gated Fusion block, proved to be structurally paramount. The removal of these modules precipitated a rapid decay in the model’s capacity to recognize spatially distant motifs. Because toxicity often arises from non-contiguous amino acid interactions folding into proximate 3D space (e.g., amphipathic helical faces), the DCNN’s expanded receptive field, combined with the gate’s selective information percolation, proved computationally indispensable.

The Cross-Attention Trajectories includes mechanism bridging 1D sequence encodings with global representational states via cross-attention served as an efficient funnel for dominant global features harvested from ESM (Figure 4). Architectures deficient in this module suffered profound instability, losing the critical linkage between localized amino-acid perturbations and their global stereochemical consequences (Table 2).

**Table 2.**
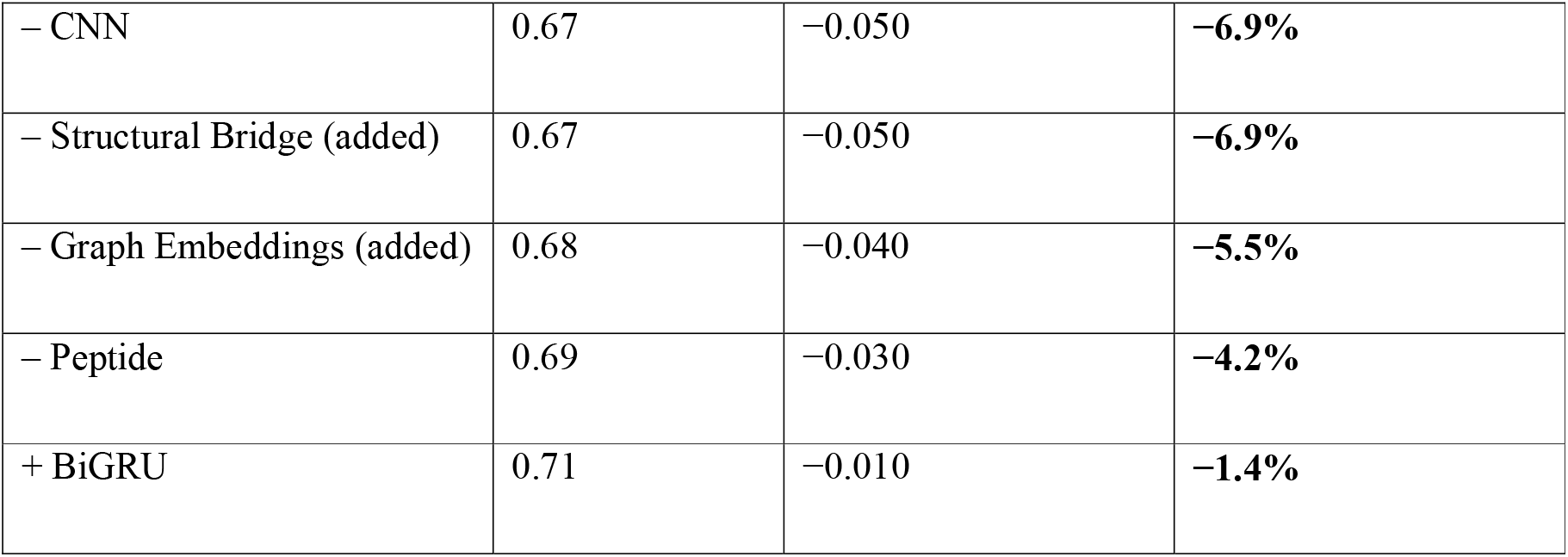
Ablation of each component and their impact on the architecture and the superior performance of the full LORA enabled architecture of GraphTox.

|  |  |  |  |
| --- | --- | --- | --- |
| – CNN | 0.67 | –0.050 | **–6.9%** |
| – Structural Bridge (added) | 0.67 | –0.050 | **–6.9%** |
| – Graph Embeddings (added) | 0.68 | –0.040 | **–5.5%** |
| – Peptide | 0.69 | –0.030 | **–4.2%** |
| + BiGRU | 0.71 | –0.010 | **–1.4%** |

**Figure 4.**
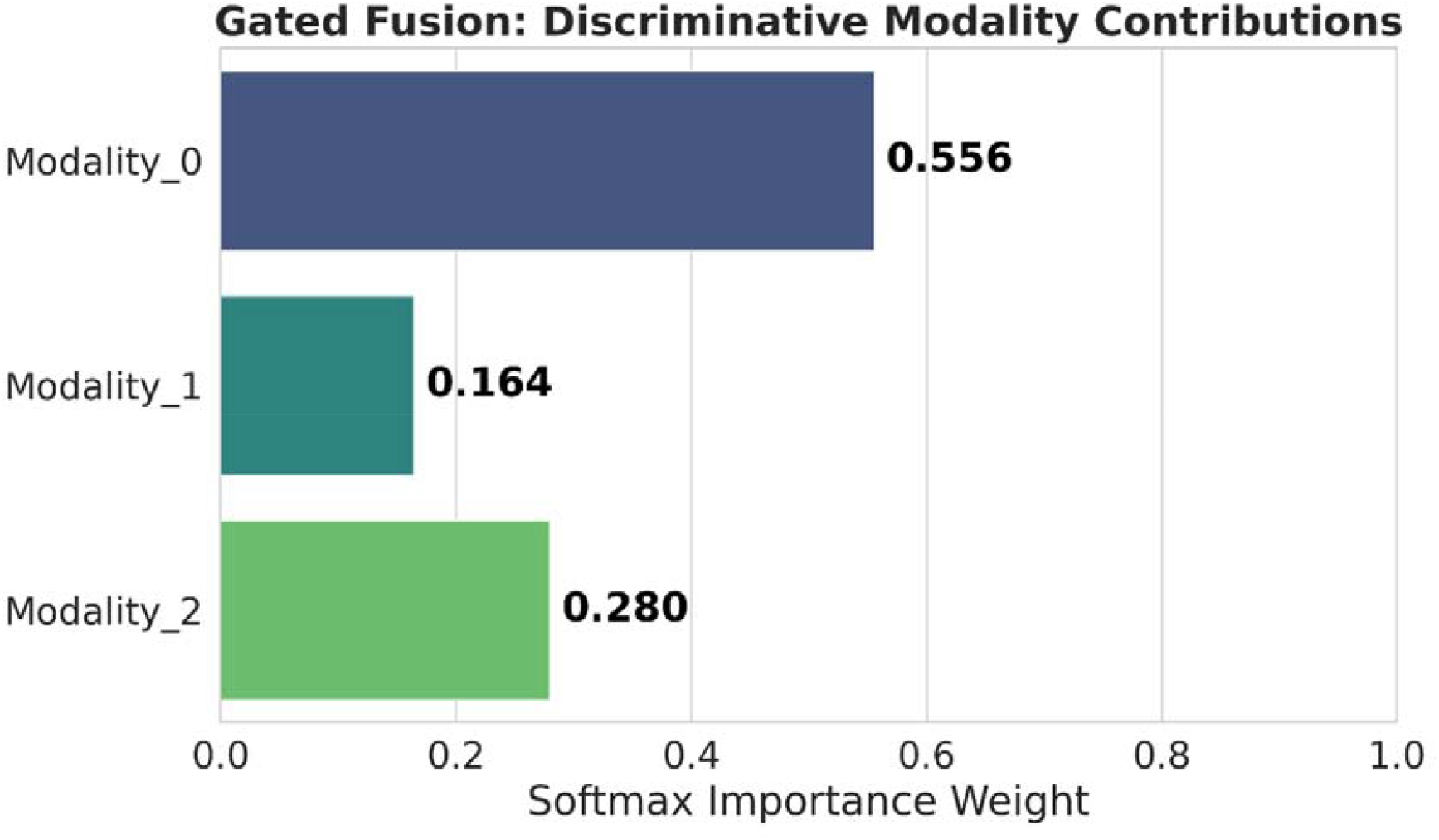
This figure shows the modality contribution of each modality where modality_0 represents UNet Sequence-Structure Bridge’, modality_1 represents ‘Global MGAE Topology’, modality_2 represents ‘Raw Peptide 1D

### 3.4 Interpretability Heuristics Unveil Robust Structural Motif Weighting Regimes

To mechanistically validate how GraphTox achieves high-fidelity toxicity discrimination, we interrogated the model’s internal representation dynamics and its biological reasoning constraints. We first visualized the evolutionary learning trajectory of peptide embeddings across sequentially critical stages of the architecture using t-SNE dimensionality reduction (Fig. 5). At the initial embedding stage following the ReLORA-perturbed ESM layer, the representations of toxic and non-toxic peptides exhibit preliminary but highly entangled clustering (tsne_esm_lora), indicating that raw sequence-level features are insufficient for robust classification. However, as these representations are parametrically projected through the pretrained Masked Graph Autoencoder (tsne_pretrained_mgae), structural geometries begin to distinctively orient the molecular topologies.

**Figure 5:**
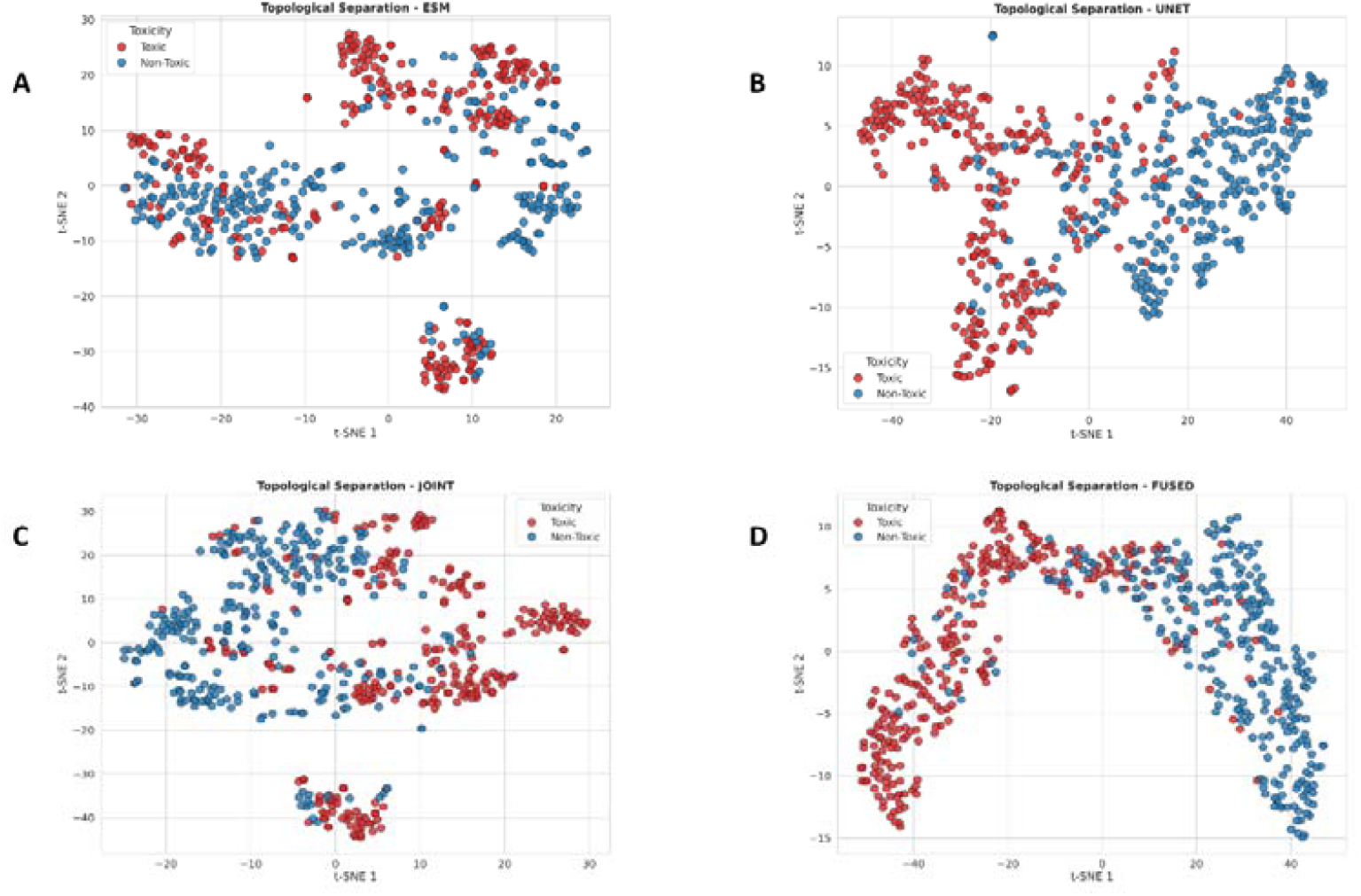
T-SNE plots from various layers 1. ESM-2 Sequence with LoRA Adaptations (tsne_esm_lora.png) LoRA-adapted ESM-2 representations remain largely intermixed, reflecting evolutionary sequence features with only weak emergent local clustering.2. Early Joint Projection Manifold (tsne_joint.png) Initial sequence–structure alignment compresses feature variance but retains substantial class overlap, indicating incomplete discriminative integration.3. Sequence-Structure U-Net Output (tsne_unet.png) Cross-attentive sequence–structure fusion induces clear manifold polarization, enabling separation of previously entangled class clusters.4. Multi-Modal Fusion Intersections (tsne_fusion_combined.png) Adaptive multi-modal fusion yields highly disentangled latent clusters, with strong class-specific segregation driven by gated feature integration.

The most profound topological partition occurs during the multimodal integration phase within the U-Net contextual sequence-structure fusion (tsne_unet) and the subsequent joint-fusion maps (tsne_fusion_combined / tsne_joint). Here, the latent manifolds undergo a distinct bifurcation. By the final classification stage (tsne_classifier), the feature distributions are decisively separated. This progressive untangling empirically validates our hypothesis: iterative, end-to-end multimodal fusion strictly exploits spatial and sequential graph attributes to construct a deterministic boundary between fundamentally similar peptide sequences.

While high predictive performance is critical, establishing the biological interpretability of these decision boundaries is paramount for downstream therapeutic design. To decode the model’s structural reasoning, we extracted deep gradient attributions and analyzed the localized motifs driving toxicity predictions. For the highest-confidence toxic classifications, GraphTox assigns heavily amplified attention weights to specific, highly localized micro-domains (Figure 6). Visual dissection of these top-ranked positive motifs consistently reveals dense functional cores characterized by hydrophobic and uniquely charged surface architectures—structural hallmarks classically associated with pore-forming or membrane-disrupting molecular behavior. Conversely, negative attribution analysis (figure 6) demonstrates that GraphTox systematically down-weights these regions in benign peptides, instead identifying stabilizing, non-reactive structural loops as biomarkers for non-toxicity. Crucially, this confirms that the architecture is not merely memorizing superficial global sequence statistics, but is actively pinpointing actionable biological micro-domains.

**Figure 6:**
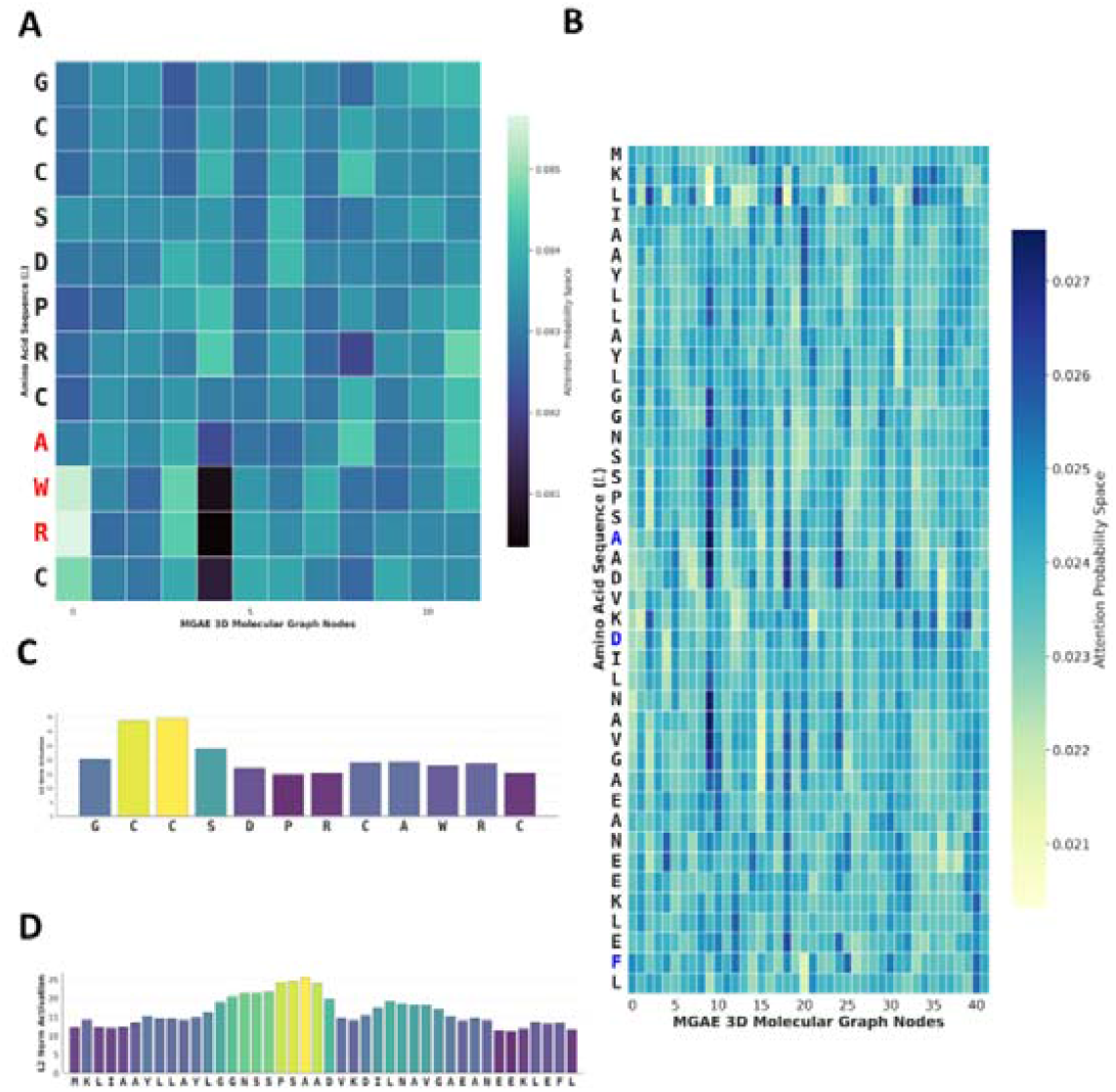
Structural attention and residue activation analysis of toxic peptide representations learned by GraphTox. (A) Attention probability heatmap illustrating residue-level interactions between amino acid positions and MGAE-derived 3D molecular graph nodes for a representative toxic peptide sequence. Residues highlighted in red (A, W, and R) exhibit distinct attention distributions, suggesting their potential contribution to toxicity-associated structural patterns. (B) Global attention probability map for a longer non negative peptide sequence showing the spatial attention learned across graph nodes and amino acid residues. Residues highlighted in blue indicate positions with elevated structural importance captured by the geometric graph transformer encoder. The attention maps demonstrate the ability of the model to capture long-range spatial dependencies and residue-specific structural interactions within peptide conformations. (C) L2 norm activation profile of residues in the short toxic peptide sequence shown in panel A. Higher activation values indicate residues contributing strongly to the learned toxicity representation, with cysteine-rich and aromatic residues exhibiting increased activation intensities. (D) Residue-wise L2 norm activation profile for the peptide sequence shown in panel B, revealing distinct peaks in activation across multiple residue positions associated with structurally informative regions. Collectively, these analyses demonstrate that GraphTox successfully identifies biologically relevant residue-level and structural interaction patterns underlying peptide toxicity through geometry-aware attention learning.

Furthermore, analyzing these interpretability heuristics elucidates the profound impact of our ReLORA fine-tuning paradigm. We mapped the internal multi-modal gradient contributions (UNet_SeqStructure_model vs. contributing_Peptide) to understand how structural awareness shifts the model’s reasoning. In the standard No-LORA baseline, the network allocated approximately 80% (0.799) of its attention to the U-Net Sequence-Structure bridge and 20% (0.200) to raw peptide features. Conversely, under the stabilized, low-rank ReLORA paradigm, the network aggressively redirected its attention, localizing over 90% (0.9037) of the attribution weight exclusively within the dense U-Net Sequence-Structure manifold and shrinking its reliance on superficial peptide features to less than 10% (0.0962). This behavioral shift provides compelling evidence that ReLORA pretraining transcends baseline sequence memorization; by stabilizing the embedding manifold, it forces the network to actively prioritize and strictly heavily weight the physically realistic, toxicologically relevant 3D spatial architectures engineered during our pretraining protocol. To mechanistically validate how GraphTox achieves high-fidelity toxicity discrimination, we interrogated the model’s internal representation dynamics and its biological reasoning constraints. To achieve this without altering the architecture’s native computational graph, we implemented a custom interpretability pipeline utilizing PyTorch forward hooks to systematically extract intermediate latent activations across critical network checkpoints. We first visualized the evolutionary learning trajectory of peptide embeddings using t-SNE dimensionality reduction (Figure 6). At the initial embedding stage—captured directly from the pooled ReLORA-perturbed ESM representations—toxic and non-toxic peptides exhibit preliminary but highly entangled clustering, indicating that raw sequence-level features are insufficient for robust classification. However, as these representations are parametrically projected through the pretrained Masked Graph Autoencoder, structural geometries begin to distinctively orient the molecular topologies. By tapping into the U-Net contextual sequence-structure fusion layers and subsequent joint-fusion mapping dimensions, we observe the most profound topological partition; the latent manifolds undergo a distinct bifurcation. By the time outputs reach the final classifier layer hook, the feature distributions are decisively separated. This progressive untangling empirically validates our hypothesis: iterative, end-to-end multimodal fusion strictly exploits spatial and sequential graph attributes to construct a deterministic boundary between fundamentally similar peptide sequences.

While high predictive performance is critical, establishing the biological interpretability of these decision boundaries is paramount for downstream therapeutic design. To decode the model’s structural reasoning, we moved beyond native forward activations by implementing an input-times-gradient (Input × Gradient) saliency mechanism. By computing the exact derivative of the predicted toxicity probability with respect to the input ReLORA-ESM embeddings—and multiplying the resulting gradient magnitude by the input representation— we robustly isolated the specific amino acid drivers of toxicity. Concurrently, we captured intermediate convolutional representations by systematically hooking into the U-Net contracting layers to extract calculations of high-resolution local motif magnitudes, and into the network’s bottleneck to identify low-resolution global domains. To explicitly track how the model maps 1D sequence queries to 3D spatial properties, we further extracted the raw cross-modal attention matrices operating deeply within the sequence-structural bridge.

Integrating these deep attribution maps revealed that for the highest-confidence toxic classifications, GraphTox assigns heavily amplified attention weights to precisely localized micro-domains (Fig. 6). Visual dissection of these top-ranked positive saliency motifs consistently reveals dense functional cores characterized by hydrophobic and uniquely charged surface architectures—hallmarks classically associated with pore-forming or membrane-disrupting molecular behavior. Conversely, negative attribution analysis systematically down-weights these regions in benign peptides, instead extracting stabilizing, non-reactive structural loops as biomarkers for non-toxicity. Crucially, the captured cross-attention mappings confirm that the architecture is not merely memorizing superficial global sequence statistics; it is actively pinpointing actionable biological micro-domains grounded continuously in 3D graph topography.

Furthermore, analyzing the numerical behavior of these internal mechanics elucidates the profound impact of our ReLORA fine-tuning paradigm. To mathematically quantify the fusion’s topological reasoning rules, we hooked into the pre-softmax gates of our multi-modal fusion layer. Mapping these internal learned parameters highlighted how deep structural awareness radically shifts the model’s decision boundary. In the standard No-LORA baseline, the network allocated approximately 80% (0.799) of its attention gate weight to the U-Net Sequence-Structure bridge and 20% (0.200) to raw peptide features. Conversely, under the stabilized, low-rank ReLORA paradigm, the network aggressively redirected its attention. It localized over 90% (0.9037) of the attribution weight exclusively within the dense U-Net topological manifold, while shrinking its reliance on superficial sequence components to less than 10% (0.0962). This behavioral shift provides compelling evidence that ReLORA pretraining transcends baseline sequence memorization; by aggressively stabilizing the embedding manifold, it forces the network to actively decode, prioritize, and strictly weight the physically realistic, toxicologically relevant 3D spatial architectures constructed during our pretraining protocol. The broader and upward-shifted distributions observed in the toxic class suggest enhanced representation learning around these residues, potentially reflecting their involvement in membrane interaction, electrostatic imbalance, structural flexibility, or residue clustering associated with toxic peptide behaviour. In contrast, non-toxic peptides display comparatively lower and more compact activation distributions, indicating weaker contribution of these motifs to non-toxic functional patterns. The statistical significance markers further validate that the observed differences are unlikely to arise from random variation, supporting the biological relevance of these amino acid motifs in discriminating toxic from non-toxic peptides (Figure 7). Overall, this analysis enhances the interpretability of the proposed model by revealing biologically meaningful residue-level patterns that contribute to peptide toxicity prediction.

**Figure 7:**
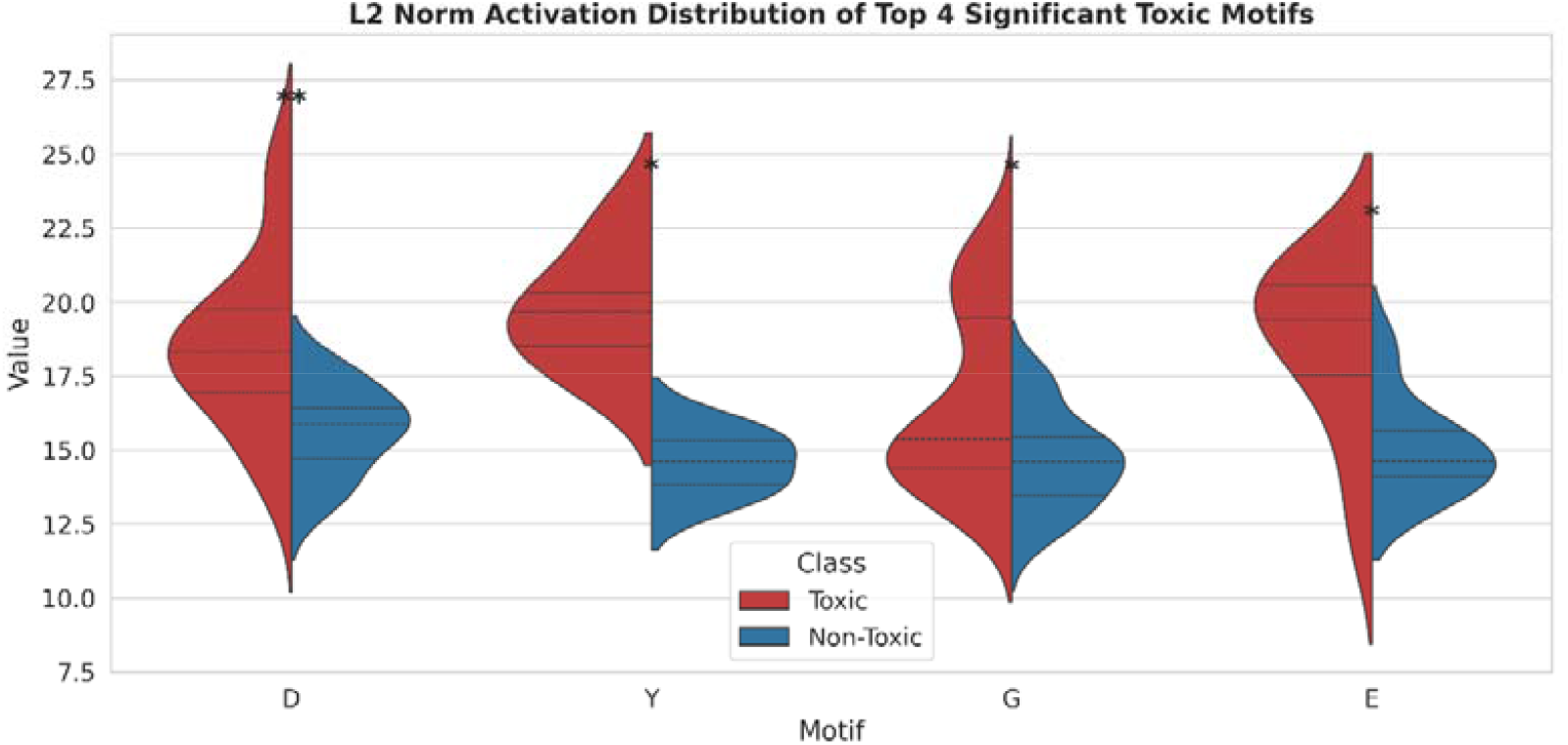
L2 norm activation distribution of the top four statistically significant toxic motifs (D, Y, G, and E) in toxic and non-toxic peptides. Split violin plots illustrate the distribution of activation values for each amino acid motif across toxic (red) and non-toxic (blue) peptide classes. The width of each violin represents the density of observations, while dashed horizontal lines indicate quartile distributions. Statistical significance between the two classes was evaluated using appropriate hypothesis testing, where asterisks denote significant differences (*p < 0.05, **p < 0.01). Toxic peptides consistently exhibit higher activation values for these motifs compared to non-toxic peptides, suggesting their strong contribution to toxicity-associated structural and biochemical patterns learned by the model.

## 4. Conclusion

Accurate prediction of peptide toxicity is critical for safe and effective development of peptide-based therapies. The conventional experimental methods for toxicity assessment are frequently expensive, laborious, and time-consuming, resulting in a high demand for credible computer prediction frameworks. Although promising performance has been shown for recent machine learning and deep learning methods, most existing methods rely heavily on sequence-derived information and are ineffective in capturing the three-dimensional structural characteristics that are critical to peptide toxicity. We describe GraphTox, a new structure-aware approach for predicting peptide toxicity that combines self-supervised geometric representation learning and hierarchical graph-based modelling. GraphTox portrays peptide structures as geometric molecular graphs and leverages a Masked Graph Autoencoder (MGAE) comprising Geometric Graph Transformer (GGT) and Equivariant Graph Neural Network (EGNN) layers to learn geometric-aware structural embeddings of peptide conformations. The proposed approach combines amino acid embeddings, spatial residue connections, and distance-aware edge representations, to capture both local residue interactions and global conformational dependencies pertinent to the peptide toxicity. Furthermore, the multi-scale U-Net architecture allows for hierarchical fusion of structural information across different resolutions, improving the model’s ability to detect spatial patterns associated with toxicity. Benchmark peptide toxicity datasets are used to show that GraphTox outperforms existing state-of-the-art approaches on many evaluation metrics. Moreover, motif activation analysis and visualisation of structural attention give biological interpretability by revealing critical residues and spatial interaction patterns contributing to toxicity prediction. These results show the significance of combining 3D structure knowledge and self-supervised geometric deep learning in peptide toxicity prediction. We propose that GraphTox provides a potent and interpretable computational framework for peptide safety assessment, and may further advance larger applications in structure-based peptide engineering, medicinal design, and biomolecular property prediction.

## CRediT authorship contribution statement

DD: Writing – review & editing, Writing – original draft, review & editing, Writing – original draft, Data curation, Validation Illustration SB: Writing – review & editing, Writing – original draft, Conceptualization, Implementation, Methodology, Data curation, Validation, Analysis, Illustration

## Acknowledgements

This work is supported by Department of Science and Technology and Biotechnology (Please Add Project Details). Additionally, Debraj (PMRF Id: 2402784) thanks the MOE, India, for the Prime Minister research fellowship.

